# Neural Representations of Task-relevant and Task-irrelevant Features of Attended Objects

**DOI:** 10.1101/2021.05.21.445168

**Authors:** Jiageng Chen, Paul S. Scotti, Emma Wu Dowd, Julie D. Golomb

## Abstract

Visual attention plays an essential role in selecting task-relevant and ignoring task-irrelevant information, for both object features and their locations. In the real world, multiple objects with multiple features are often simultaneously present in a scene. When spatial attention selects an object, how are the task-relevant and task-irrelevant features represented in the brain? Previous literature has shown conflicting results on whether and how irrelevant features are represented in visual cortex. In an fMRI task, we used a modified inverted encoding model (IEM, e.g., Sprague & Serences, 2015) to test whether we can reconstruct the task-relevant and task-irrelevant features of spatially attended objects in a multi-feature (color + orientation), multi-item display. Subjects were briefly shown an array of three colored, oriented gratings. Subjects were instructed as to which feature (color or orientation) was relevant before each block, and on each trial were asked to report the task-relevant feature of the object that appeared at a spatially pre-cued location, using a continuous color or orientation wheel. By applying the IEM, we achieved reliable feature reconstructions for the task-relevant features of the attended object from visual ROIs (V1 and V4v) and Intraparietal sulcus. Preliminary searchlight analyses showed that task-irrelevant features of attended objects could be reconstructed from activity in some intraparietal areas, but the reconstructions were much weaker and less reliable compared with task-relevant features. These results suggest that both relevant and irrelevant features may be represented in visual and parietal cortex but in different forms. Our method provides potential tools to noninvasively measure unattended feature representations and probe the extent to which spatial attention acts as a “glue” to bind task-relevant and task-irrelevant features.

## Introduction

The visual world is complex. It usually contains far more objects and features than humans can effectively process at a time. As we have limited cognitive resources, we are able to deploy attention resources strategically and prioritize the most relevant objects and features based on the task demands. When spatial attention selects an object, how are the task relevant and task irrelevant features represented in the brain? There has been much of neurophysiological evidence showing that frontal, parietal and sensory cortices exhibit feature selectivity for task-relevant stimuli (Bichot, et al., 1996; Ibos & Freedman, 2014; Toth & Assad, 2002; Wurtz & Mohler, 1976; Moran & Desimone, 1985). Recently, fMRI studies have shown that using multivariate pattern analysis (MVPA) and inverted encoding models (IEM), task-relevant features can be decoded and reconstructed in variety of brain regions (Albers, et al., 2013; Bettencourt& Xu, 2016; Brouwer & Heeger, 2009, 2011; Christophel et al., 2012; Ester et al., 2015; Harrison & Tong, 2009; Serences et al., 2009; Yu & Shim, 2017).

However, a few important questions remain untested. First, most of the prior studies have presented either single items at the center of the screen (e.g. Brouwer & Heeger, 2009), or 2 items of a single feature (e.g., 2 gabors of different orientations: Ester et al., 2015). Real world conditions often contain multiple objects of multiple features. Second, it remains unclear how spatial and feature attention interact in these representations. For example, representations are enhanced for spatially attended objects (Desimone & Duncan, 1995; Somers, et al., 1999), and it is possible to reconstruct the task-relevant feature of an object (Yu & Shim, 2017), but less clear is whether and how our brain represents *task-irrelevant* features, especially the task-irrelevant features of a spatially cued or attended object. Behavioral studies suggest that task-irrelevant features of a cued object might still be processed at certain levels. For example, studies of object-based attention found that attention could spread along the cued object and facilitate the processing of other features within the object (Duncan, 1984; Egly et al., 1994; O’Craven et al., 1999). But behavioral studies are limited in their ability to study truly task-irrelevant information. Neuroimaging may be better suited, but previous studies have shown mixed results: In the visual working memory literature, most studies have not found evidence supporting the active representation of unattended features (LaRocque et al., 2017; Lewis-Peacock et al., 2012). But these studies often tested unattended features of *uncued* stimuli, which are less likely to be processed overall. Literature have suggested that the unattended contents of working memory cannot be directly decoded but are stored in an activity-silent form (Stokes, 2015; Wolff et al., 2017). Contrarily, other studies have found that intraparietal areas and the frontal eye fields maintained a low-resolution representation of unattended item (Christophel et al., 2018). On the other hand, in the object-processing literature, studies have shown that task-irrelevant features of an attended object activate category-selective brain areas to an increased extent (Xu, 2010).

In the current study, we tested whether we could reconstruct the task-relevant and task-irrelevant features of spatially attended objects in a multi-feature (color + orientation), multi-item display using fMRI. Participants were presented with an array of three colored, oriented gratings and were asked to report either the color or orientation (i.e., the task-relevant feature) of the object that appeared at a spatially pre-cued location. By applying a modified IEM, we asked if and where we could reconstruct the task-relevant and task-irrelevant features of the cued object. We chose a perception task over presenting a long memory delay because previous studies suggested that information about task-irrelevant features could be short-lived (Xu, 2010).

Beyond our first goal, we also aimed to use this technique to test how the reconstruction of task-relevant and task-irrelevant features change when spatial attention is not stable. Because the visual environment is dynamic and our behavioral goals may also change across time, our attention is not static and often shifts between objects and features. Previous literature has demonstrated that shifts of spatial attention can cause systematic behavioral errors, for example, reporting a previously attended stimulus or showing systematic behavioral bias (Chen et al., 2019; Golomb, 2015; Golomb et al., 2014). However, we haven’t fully understood the neural mechanisms of those behavioral errors and if/how a shift of attention alters the neural representations to cause the errors. In the current study, we included a mix of Hold spatial attention and Shift spatial attention trials; on the latter participants shifted spatial attention just before the stimulus array was presented. We predicted that participants might exhibit systematic behavioral errors (swap errors) on some of the Shift trials, and we sought to test the neural reconstructions of those trials on which participants made swapping errors and build links between behavioral performance and neural activity.

## Method

### Participants

9 students (ages 18-33, mean age:24.1; 5 females) from The Ohio State University participated in the experiment for monetary compensation ($10/hour for behavioral prescreening and $15/hour for fMRI sessions). All participants reported having normal color vision and normal (or corrected-to-normal) visual acuity. Written informed consent was obtained for all participants and study protocols were approved by The Ohio State University Behavioral and Social Sciences Institutional Review Board.

### Stimuli and design

All stimuli were generated using MATLAB (MathWorks, Natick, MA) and the Psychophysics Toolbox (Brainard, 1997;Kleiner, Brainard, & Pelli, 2007; Pelli, 1997) on a Macbook Pro laptop.

Stimuli were projected onto a screen behind the scanner core and participants viewed the stimuli via a mirror at 45° above the head coil. The viewing distance was 74 cm. Eye position was monitored using an Eyelink 1000 system.

The stimulus array contained three colored and oriented gratings on a black background, located at 12 o’clock, 4 o’clock and 8 o’clock positions on the screen with 5° radius from central fixation. The diameter of each grating was 5°. The colors of gratings were picked from a set of color values along a color wheel in CIEL*a*b* color space. The color wheel was centered at (L*=70, a*=20, b*=38) with a radius of 60. We first defined a set of 8 base colors along the color wheel, such that adjacent base colors were spaced 45° apart in circular color space. For each trial, we selected the target color from one of the 8 base colors, and then added a positive or negative 1-10° jitter. The other two non-target colors were then selected such that the three colors were each exactly 120° apart. Similarly, for orientation we defined 8 base orientations with increments of 22.5°. For each trial, we selected the target orientation from the base set, and then added a positive or negative 1-5° jitter. The other two non-target orientations were then picked such that all three orientations were 60° apart.

The task schematic is shown in Figure 1. Trials were blocked into Attend Color and Attend Orientation trials. Each trial started with a fixation cross (size: 0.5° line thickness, white) at the center of the screen. After 700ms, three circle outlines were displayed at the stimulus locations for 200ms. Two outlines were placeholders (0.1° line thickness, gray) and one was the spatial cue (0.3° line thickness, white). Participants were instructed to covertly attend to the spatial cue location while maintaining eye fixation at the screen center.

**Figure 1.**
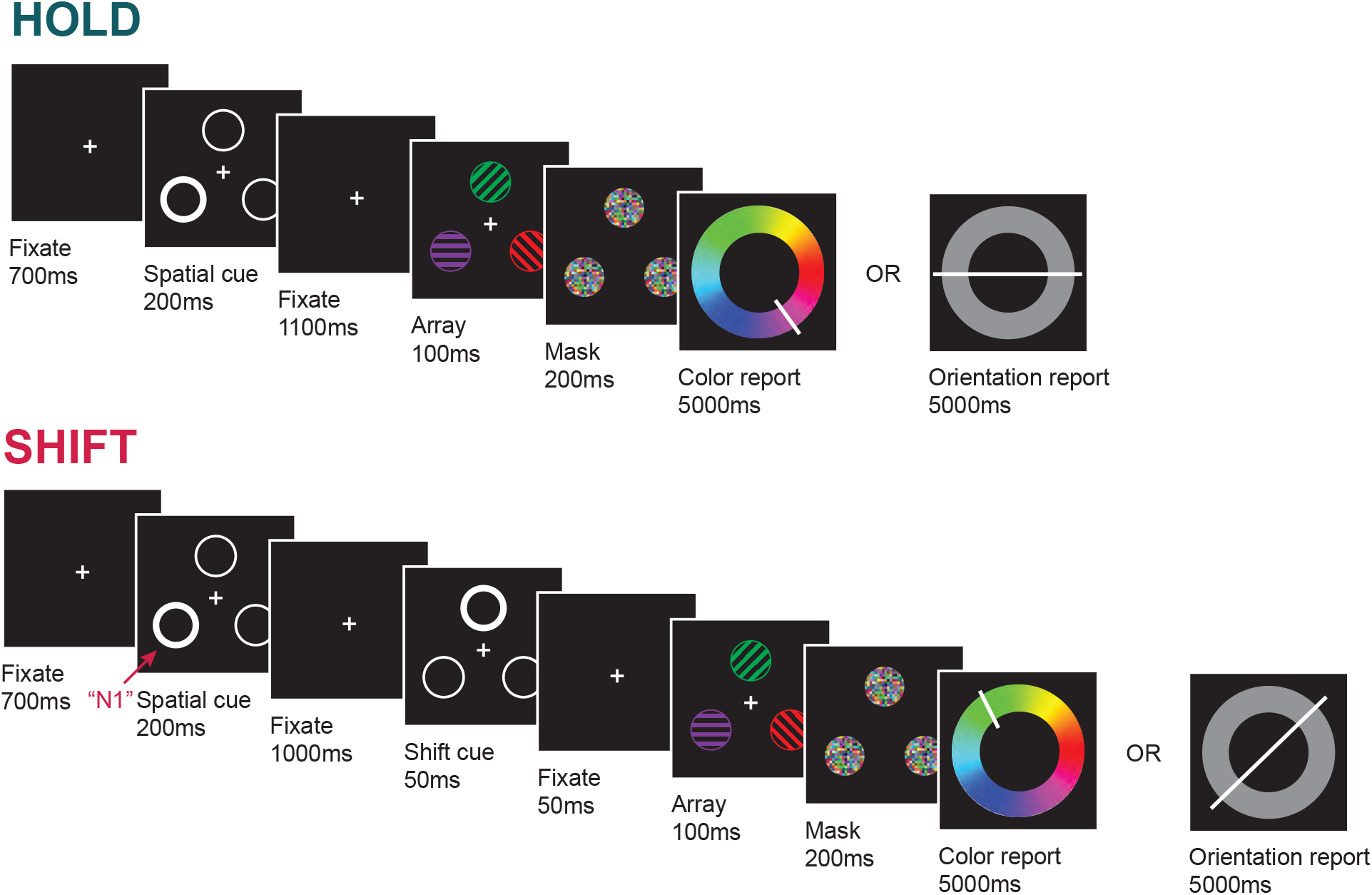
fMRI task schematic. In hold trials, the spatial cue was presented for 200ms. After a 1100ms fixation interval, the stimulus array was presented briefly for 100ms, followed by a 200ms mask and a 5000ms response screen. Participants pressed keys in the button box to move the white line to match the color (Color run) or orientation (Orientation run) of the gratings that appeared at the cued location. The trial sequence of shift trials was the same as the hold trials, except a shift cue was displayed 1000ms after the initial spatial cue disappeared. Participants were instructed to report the color/orientation of the grating that appeared at the shift cue location.

In half of trials (Hold trials), after a 1100ms fixation interval, the stimulus array was displayed briefly for 100ms, followed by a 200ms mask and a color or orientation report screen (depending on the task). Participants were instructed to report the color (Color run) or orientation (Orientation run) of the grating that appeared at the location of the spatial cue (the “target”), using a modified continuous report technique. Participants controlled four buttons to adjust the color/orientation until it matched the target feature (a white line was superimposed on the color or orientation wheel, and the 4 buttons allowed for both coarse and fine adjustments in clockwise and counterclockwise directions). Participants had 5 seconds to complete their response, and the final color/orientation at the end of the 5 second window was taken as the report.

In the other half of trials (Shift trials), 1000ms after the initial spatial cue disappeared, a second spatial cue appeared for 50ms. The second spatial cue was always at a different location from the first spatial cue. Participants were instructed to shift covert attention to the new location if a second cue appeared, and to report the color/orientation of the item appearing at that most recently cued location. The stimulus array appeared 50ms after the shift cue disappeared, and the remainder of the trial was identical to Hold trials. The timing of the shift cue (100ms before the stimulus array) was selected based on prior behavioral studies to optimize the likelihood that Shift trials would reflect of mixture of trials on which attention successfully shifted to the new location and trials on which attention was still stuck at the originally cued location at the time of stimulus presentation (Dowd & Golomb, 2019; Golomb et al., 2014).

Each trial lasted 7.3 seconds, and each run had 40 trials, with 20 Hold and 20 Shift trials intermixed. Trial start times were jittered and synced to the fMRI scanner TRs, with stimulus onset asynchrony randomly chosen from 8, 10, or 12s in proportions of 50%, 35%, and 15%, respectively. Participants either reported color or orientation for the cued stimulus in each run; the feature that participants were asked to report remained the same through the entire run. Color runs and Orientation runs alternated through a 2-hour scanning session. Each participant completed multiple scanning sessions, for a total of at least 22 runs (22 to 40 runs). In addition, each participant also completed 2 standard retinotopic mapping runs.

### fMRI acquisition

Each participant was scanned in two 2-hour fMRI sessions (one participant in four 2-hour sessions). All scanning was carried out on a 3T Siemens Prisma scanner using a 32-channel head coil at the Center for Cognitive and Behavioral Brain Imaging (CCBBI) at The Ohio State University. Functional data were acquired with a T2-weighted gradient-echo sequence (TR = 2000 ms, TE = 28 ms, 72° flip angle). Slices were oriented to maximize coverage of occipital lobe and posterior parietal lobe (33 slices, 2*2*2mm voxels, 0% gap). A high-resolution (1*1*1 mm) MPRAGE anatomical scan was acquired before the functional scans in each fMRI session.

### Behavioral analyses

We calculated the feature report error for each trial by subtracting the correct feature value from the reported feature value, such that an error of 0° is a perfect response. For color, the error range is from -180 to 180; for orientation, the error ranges from -90 to 90. For illustration purposes, we aligned the direction of report errors in the Shift condition so that a positively signed error means the reported feature was shifted in feature space towards the feature of the item at the initially cued location; a negative report error means the reported feature is farther away in color space from the feature of the initially cued item. On hold trials, an analogous mock alignment was performed, where the sign of report errors was randomly flipped on half of the trials (Chen et al., 2019).

We also fitted the distribution of errors for each condition (Hold and Shift for Color and Orientation trials) using a probabilistic mixture model. The model assumes the distribution of report error comes from several sources (Formula 1): one von Mises distribution (ϕ) accounting for the probability to correctly report the target feature, with a flexible mean (μ) allowing the model to capture any systematic bias from the target feature and a concentration parameter (κ) reflecting precision; two von Mises distributions accounting for the probabilities (*β*_1_, *β*_2_) to misreport the feature of one of the two non-target items instead (where *β*_1_ is the item at the originally cued location); and one uniform distribution accounting for the probability (γ) of random guessing.

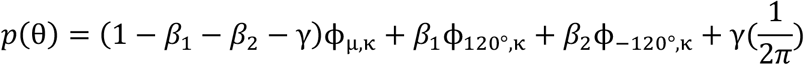

For each participant and each condition, we fit the model by applying Markov chain Monte Carlo using MemToolbox (Suchow et al., 2013). The best-fitting parameters (maximum likelihood estimate) were submitted to the paired-sample test and repeated ANOVAs. We compared the parameter differences between conditions. We also tested whether there were feature distortions in each condition by comparing the mean of the von Mises distribution (μ) to 0.

### fMRI preprocessing

fMRI data preprocessing was conducted using SPM12 and custom MATLAB code. The functional data were corrected for slice acquisition times and spatially realigned to the first volume of the first session to account for head motions. A 128-s temporal high-pass filter was applied to remove the low frequency noise. Structural images from different fMRI sessions were averaged and coregistered with functional images. For univariate (Stimulus Array > Fixation) contrast only, we spatially smoothed the data with a Gaussian kernel of 4mm FWHM. For other multivariate analyses (IEM or decoding) we kept data unsmoothed and within the native space for each participant. For data analyses and visualization purposes, we subsequently smoothed and normalized searchlight results into standard MNI space. See *Retinotopic mapping, Inverted encoding model analyses* and *Searchlight* for details.

### Retinotopic mapping

We applied a standard retinotopic mapping localizer to define areas V1 and V4v (Sereno et al., 1995). Each participant completed one pair of retinotopic mapping runs to measure the polar angle maps. A 60° high contrast checkerboard wedge rotated around a central fixation dot on the screen and flickered at 4 Hz. The eccentricity of the wedge spanned from the central 1.6° foveal region to 16° peripheral region. The wedge rotated either clockwise or counterclockwise for 7 cycles in each run and each cycle lasted 24 seconds. Participants were instructed to maintain fixation at the center of the screen through each run and press a button whenever the fixation dot flashed.

The functional data from retinotopic mapping runs were preprocessed and analyzed using BrainVoyager QX, Freesurfer and custom MATLAB code. The resulting polar angle maps were projected onto a flatten surface map in BrainVoyager QX. The visual ROIs were drawn by hand followed by standard procedures. Each ROI was then constrained by univariate contrast of (Stimulus Array > Fixation) to select voxels that showed significant activation during the task.

### Regions of interest

Regions of interest were defined individually for each participant based on functional (V1, V4v) and anatomical (IPS) localizers. We chose V1 as the visual ROI for orientation and V4v as the visual ROI for color. We also defined an ROI in Intraparietal sulcus (IPS) from the Destrieux atlas ((Destrieux, Fischl, Dale & Halgren, 2010)) in Freesurfer (parcel labelled “S_intrapariet_and_P_trans”). For exploratory analyses, we also defined multiple additional parietal and occipital ROIs from the Destrieux atlas in Freesurfer using labels “G_parietal_sup” (Superior parietal lobule), “G_pariet_inf_Angular” (Angular gyrus), “S_oc_sup&transversal” (Superior occipital sulcus and transverse occipital sulcus), and “G_occipital_sup” (Superior occipital gyrus). We combined left and right hemispheres for all ROIs.

### Searchlight analyses

In addition to analyzing data from the ROIs listed above, exploratory searchlight analyses were also conducted to search for clusters that showed significant reconstructions for the task-relevant or task-irrelevant features. We defined a sphere ROI, with radius of 10mm (∼390 voxels), and iteratively moved it across individual brains. For each iteration, we followed the same IEM procedure described below (See *Inverted encoding model analyses*). We generated four searchlight maps for each participant: reconstructing task-relevant color, task-relevant orientation, task-irrelevant color, and task-irrelevant orientation. To better average across participants and visualize the result, the searchlight maps were normalized and spatially smoothed with an 4mm FWHM kernel. The significance level was estimated with a permutation procedure (See *Permutation test*).

### Inverted encoding model analyses

We applied a modified inverted encoding model (IEM) approach to reconstruct color and orientation information of the attended stimulus. To obtain single-trial neural activations for IEM, we modified a commonly used single-trial general linear model (GLM) approach (Mumford et al., 2012) to improve the model sensitivity and account for our large number of trials. Specifically, we conducted 40 GLMs per subject, where each GLM includes one regressor per run for *one* of the 40 trials in that run and one regressor per run for all the other remaining trials in that run. In this way, across the 40 GLMs, each trial in the experiment has an estimated single-trial beta weight.

The beta weights for each trial (of a given condition) and each voxel (within a given ROI or searchlight sphere) were extracted as the input for the IEM analyses. We separately analyzed color runs and orientation runs, and *Hold* and *Shift* trials. We applied a leave-one-run-out cross validation procedure to avoid over-fitting. To reconstruct the attended color, we used all-but-one color runs as the training set and tested the left-out color run. We iterated the procedure until all color runs served as the test run. The reconstruction of attended orientation followed the same process. To reconstruct the *unattended* color, we used all-but-one *orientation* runs as the training set (but color value as the training label) and tested the left-out orientation run.

We followed a standard IEM procedure (Sprague & Serences, 2015), with the addition of iterative shifting of the basis set (Scotti et al., in prep). This analysis assumes the neural activation patterns for each voxel reflect the linear sum of a basis set of 8 different hypothesized feature channels. Each channel has a response function modeled as the half sinusoids raised to the 7^th^ power and centered at the one of the 8-base color/orientation we defined (See *Stimulus and Design*). We conduct the analysis below iteratively, shifting the center of the basis set by 1 degree on each iteration to obtain smoother and more accurate reconstructions (Scotti et al., in prep).

Because we assume a linear relationship between the voxel activation and channel response, we model:

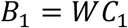

where B_1_ (m voxels * n trials) is the beta weights for each voxel, W (m voxels * k channels) is the weight matrix that characterizes a linear mapping from hypothesized channel space to voxel space, and C_1_ (k channels * n trials) is the feature channel response for the training set.

We estimate the weight matrix W by applying least-square approach using data from the training set:

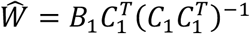

Next, we invert the model to determine the estimated feature channel response 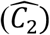 for the test set:

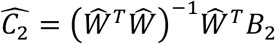

We get one feature channel response as the reconstruction function for each trial. The predicted feature of each trial was best estimated by the peak of the feature channel response. To average the reconstructed feature across trials, the estimated feature channels and the predicted features were circularly shifted so that the correct feature was at 0°.

To evaluate the reconstruction quality, we used a *reconstruction fidelity* measure (RF, Sprague et al., 2016), calculated as the vector mean of the reconstruction function:

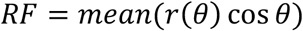

Where *θ* is the polar angle across the entire feature space (−180° to 180°) and *r*(*θ*) is the reconstruction function.

### Permutation tests

The significance of reconstruction fidelity was calculated with a permutation procedure. We followed the same analyses procedure as described above in *Inverted encoding model analyses*, but randomly shuffled each trial’s training label. We repeated the process 1000 times to create a null distribution of RF. We then compared the true RF calculated for that condition to the null distribution. The p value is the fraction of samples in the null distribution that is larger than the true RF.

### Linking neural reconstructions to behavioral errors

We also tested whether the neural reconstructions tracked participants’ behavioral reports. First, we tested whether the neural reconstructions reflected when participants reported an entirely wrong item (behavioral “Swap errors”). As a coarse approach, we evenly split the feature space into three parts and classified each trial as one of three trial types: We defined a trial as “a target report trial” if the behavioral report error was within ± 60° in color space or ± 30° in orientation space of the correct feature. If the reported color was instead within ± 60° of one of the two non-target items (± 30° for orientation), we classified the trial as “a positive swap trial (b1 trial)” or “a negative swap trial (b2 trial)”. For Hold trials, the positive and negative swap items were randomly chosen between the two non-target items. For Shift trials, we aligned the errors so that reporting the item at the originally cued location was classified as a “b1 trial”, and reporting the other non-target item was classified as a “b2 trial”. We conducted a set of IEM analyses using the approach described above, but separately for target report trials and swap trials. For the swap trials, we calculated RF as if the “correct” response was the swapped item (i.e., “swap-RF”). A significant swap-RF indicated the reconstructions successfully track the reported feature, although participants made huge swap errors in those trials. Second, we directly tested the links between the reconstruction quality and behavioral performance. We calculated the Pearson’s correlation between each trial’s reconstruction fidelity and the absolute report error across all participants. Better RF represents better reconstruction quality. We predicted better reconstruction quality is associated with better behavioral report performance. To increase our power, we combined hold trials and shift trials (5040 total trials for orientation and 5120 total trials for color).

## Results

### Participants had good behavioral performance in reporting task-relevant features

The main emphasis of this current study is the neural reconstructions (reported below). However, to verify that the paradigm was working properly and characterize the behavioral data before trying to link them with the neural reconstructions, we first report an analysis of the behavioral data. The error distributions from different conditions are shown in Figure 2a. Overall, participants reported task-relevant features rather accurately and the error distributions were centered around 0°. To further quantify the response errors, we fitted each participant’s error distribution with a mixture model (see *Analyses*).

**Figure 2.**
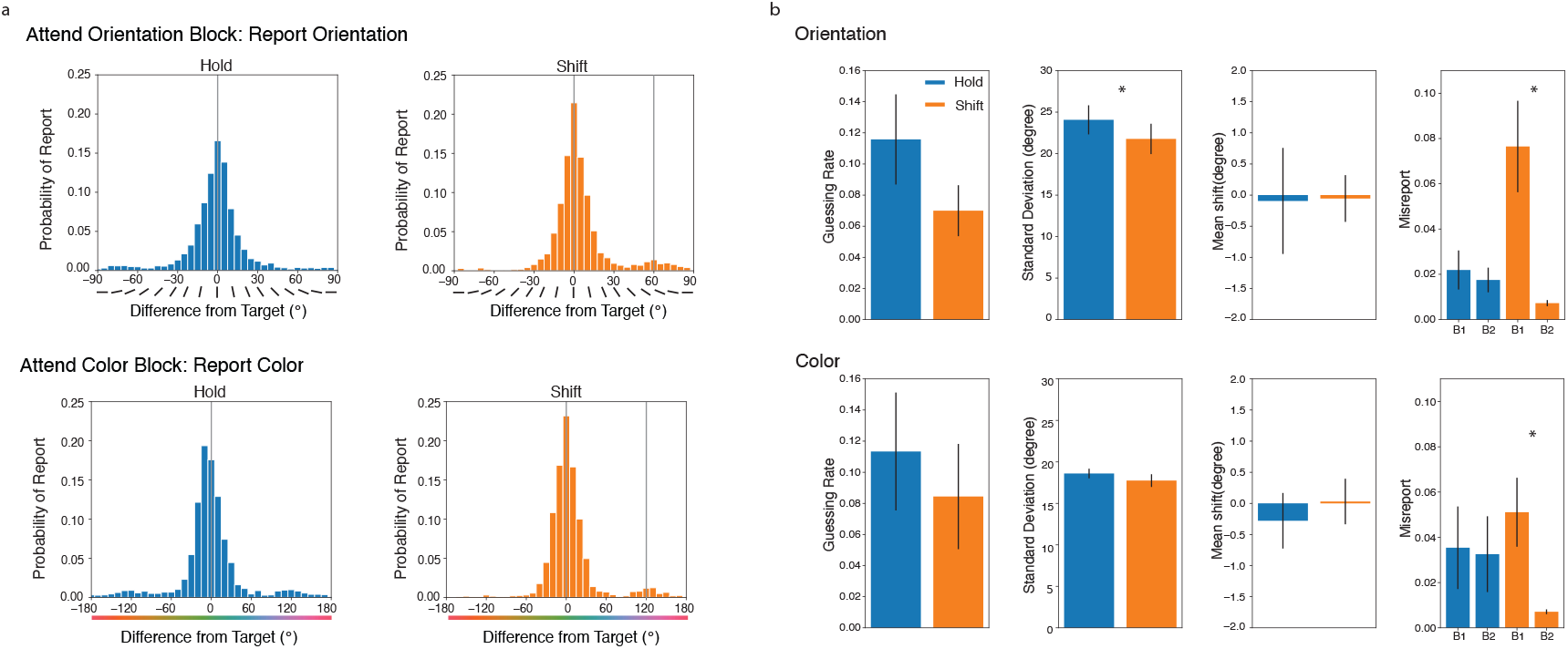
Behavioral Results. a) The error distributions from different conditions. The behavioral errors for both color and orientation reports were relatively small. b) Probabilistic mixture model results from different conditions. For shift trials, participants were more likely to report an initially cued feature (b1) than the control feature (b2) for both color and orientation runs.

In orientation runs, the probability of reporting the target (pT = 1 - Guessing rate - Swap errors) was quite high, for both hold and shift trials (Mean probability to report target: 84.48% for hold trials, 84.63% for shift trials) showing that participants correctly reported the task-relevant features for most trials. Next, we conducted a series of paired-sample t-tests and repeated measures ANOVA to compare the behavioral performance of hold trials and shift trials (Figure 2b). Overall, participants’ response in shift trials were slightly more accurate than hold trials with a lower guessing rate and higher precision (Guessing rate: t(8) = 2.181, p = 0.061; Standard deviation (inverse of precision): t(8) = 3.151, p =0.014). But compared with hold trials, participants made more swap errors to the initially cued item than the control item in shift trials: We ran a two-way repeated measures ANOVA comparing the factors of trial types (hold vs shift) and misreport item (b1 vs b2). We found a significant main effect of misreport item (F(1,8)=10.640, p=0.011), which was qualified by a significant trial type * misreport interaction (F(1,8)=10.228, p=0.013), indicating different effect of trial type on the misreport item. In hold trials, the probabilities of reporting each of the two uncued items were not significantly different from each other (t(8)=1.067, p=0.317). However, in shift trials, participants were more likely to report the feature of the initially cued item than the control item (t(8)=3.289, p=0.011), consistent with prior behavioral findings (Chen et al., 2019; Dowd & Golomb, 2019; Golomb et al., 2014). Last, we wondered if the shift cue caused any systematic feature bias. We did one-sample t-tests to compare the bias parameter (μ) to 0 for both hold and shift trials. We didn’t find evidence of feature bias in either type of in orientation report blocks (t(8) = 0.040, p = 0.969).

Similar to orientation runs, participants had good performance in color runs overall. The probability of reporting the target was high in both hold and shift trials (Mean probability to report target: 81.87% for hold trials, 85.75% for shift trials). We then compared the behavioral performance between hold and shift trials (Figure 2b). Both guessing rate and standard deviation of hold and shift trials were not significantly different (Guessing Rate: t(8)=1.801, p=0.108; Standard deviation: t(8)=1.092, p=0.307). Next, we tested whether participants were more likely to report the initially cued color in shift trials. The two-way repeated measure ANOVA again revealed a significant main effect of misreport item (F(1,8)=8.605, p=0.022) and an interaction between trial type and misreport item (F(1,8)=6.075, p=0.039). Participants were more likely to report the color of the initially cued item over the control item in shift trials (t(8)=2.730, p=0.026) but had no preference between control items in hold trials (t(8)=0.719, p=0.492). As with orientation, we did not find any feature bias for color reports in either hold or shift trials (t(8)=0.517, p=0.619).

### We can reliably reconstruct the *task-relevant* features of the attended items from neural activity patterns

Using IEM, we first asked if we could reconstruct the *task-relevant* colors and orientations of the attended target item from the patterns of neural activity in early visual and IPS ROIs. We calculated the reconstruction fidelity in each ROI and compared it with chance level. The reconstruction results are shown in Figure 3. In V1, the orientation reconstruction fidelity was significantly above chance in both hold (RF=0.163, p<0.001) and shift (RF=0.164, p<0.001) trials. The color reconstruction fidelity was marginally above chance in hold (RF=0.095, p=0.059) trials and significantly above chance in shift (RF=0.027, p=0.001) trials. In V4, the orientation reconstruction fidelity was significantly above chance in hold (RF=0.116, p=0.030) and shift (RF=0.157, p<0.001) trials. The color reconstruction fidelity was also significantly above chance in hold (RF=0.086, p=0.041) and shift (RF=0.087, p=0.043) trials. In IPS, we found reliable reconstruction fidelities in hold and shift trials in both color reporting blocks and orientation reporting blocks (Orientation: RF_hold=0.020, p=0.020; RF_shift=0.025, p=0.010; Color: RF_hold=0.027, p=0.001; RF_shift=0.036, p=0.001). Among all ROIs, the reconstruction fidelity of hold versus shift trials were not significantly different. In summary, the task-relevant features of the attended item could be reliably reconstructed in visual ROIs and IPS, even with three, multi-feature items in the display.

**Figure 3.**
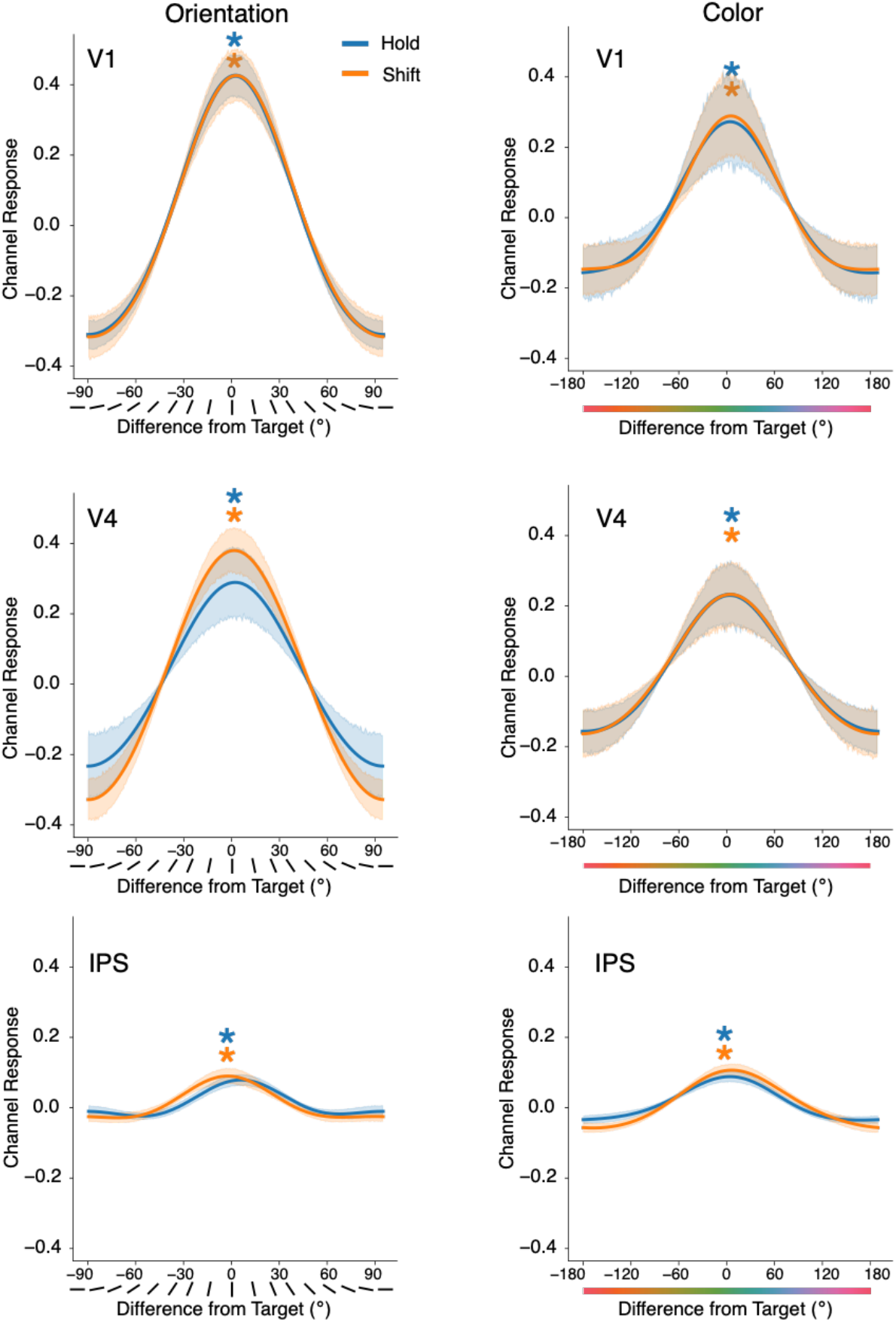
Feature reconstructions of task-relevant features of the attended item. We achieved reliable reconstructions of task-relevant features in early visual ROIs and IPS for both color and orientation runs. We did not observer any differences between hold and shift trials.

### Neural reconstructions of the task-relevant features track behavioral performance

Since we can reliably reconstruct the task-relevant features of the attended item, we next tested whether the reconstruction quality of the task-relevant features linked to participants’ behavioral reports. As we observed significant swap errors in shift trials in behavioral analyses, we asked whether the reconstructions of the task-relevant features in those trials reflect those swap errors. Analyzing the subset of Shift trials on which participants made a large behavioral report error around the initially attended feature, we found that reconstructions in V1 and V4 similarly reflected the misreported (swapped) feature (Figure 4). To better quantify the reconstructions on these trials, we calculated the swap-RF for those trials which participants made swap errors (See *Analyses*). For orientation runs, when participants reported the previous attended orientation (“b1 trials”), the swap-RF was significantly above 0 in both early visual ROIs (swap-RF_V1 = 0.157, p<0.001; swap-RF_V4: 0.137, p<0.001), indicating the reconstruction in early visual ROIs successfully tracked swap errors. For color runs, the effect was in the same direction numerically but weaker. When participants reported the previous attended color (“b1 trials”), the swap-RF of V1 and V4 were both above 0 but did not reach significance level (swap-RF_V1 = 0.064, p=0.092; swap-RF_V4: 0.063, p=0.069). On the contrary, we did not find any evidence showing IPS reconstruction could track swap errors in either orientation runs or color runs (orientation: swap-RF_IPS = -0.016, p=0.625; color: swap-RF_IPS = -0.025, p=0.236). We note that although these results are quite strong in the early visual ROIs, there is a possible alternative explanation limiting the impact of these findings (see Discussion).

**Figure 4.**
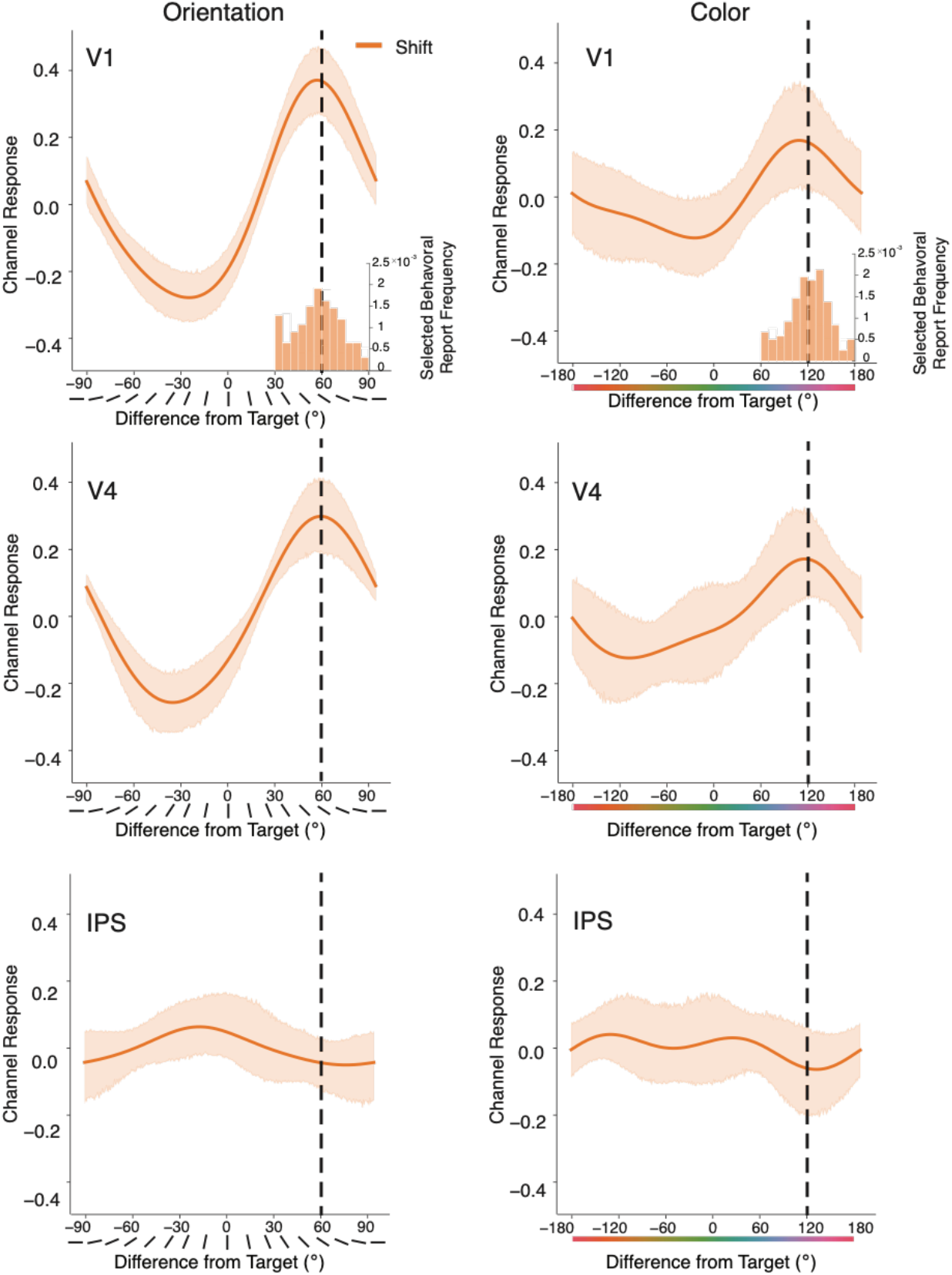
Neural reconstructions of task-relevant features when participants made large behavioral errors. Neural reconstructions of task-relevant features when participants made large behavioral errors (inset histograms) mis-reporting the initially attended feature in shift trials. Neural reconstruction of both V1 and V4 tracked the swap errors of task-relevant feature while IPS did not show any feature selectivity to the misreported feature.

In addition to the neural representations tracking these large swap errors on shift trials, we asked more generally whether the neural reconstructions were linked with behavioral reports on each trial. We calculated the Pearson’s correlation between the neural RF and behavioral absolute report error at the trial-by-trial level for both hold and shift trials (Figure 5). We observed strong negative correlations in V1 for both colors and orientations: larger behavioral errors were associated with worse neural RF (r_orientation = -0.367, p<0.001, r_color = -0.150, p<0.001). Similar effects were observed both in V4 and IPS, although the effect in IPS was weaker than early visual areas (V4: r_orientation = -0.278, p<0.001, r_color = -0.130, p<0.001; IPS: r_orientation = -0.033, p=0.024, r_color = -0.034, p=0.016). Overall, across our priori ROIs, better neural RFs were associated with smaller behavioral report errors.

**Figure 5.**
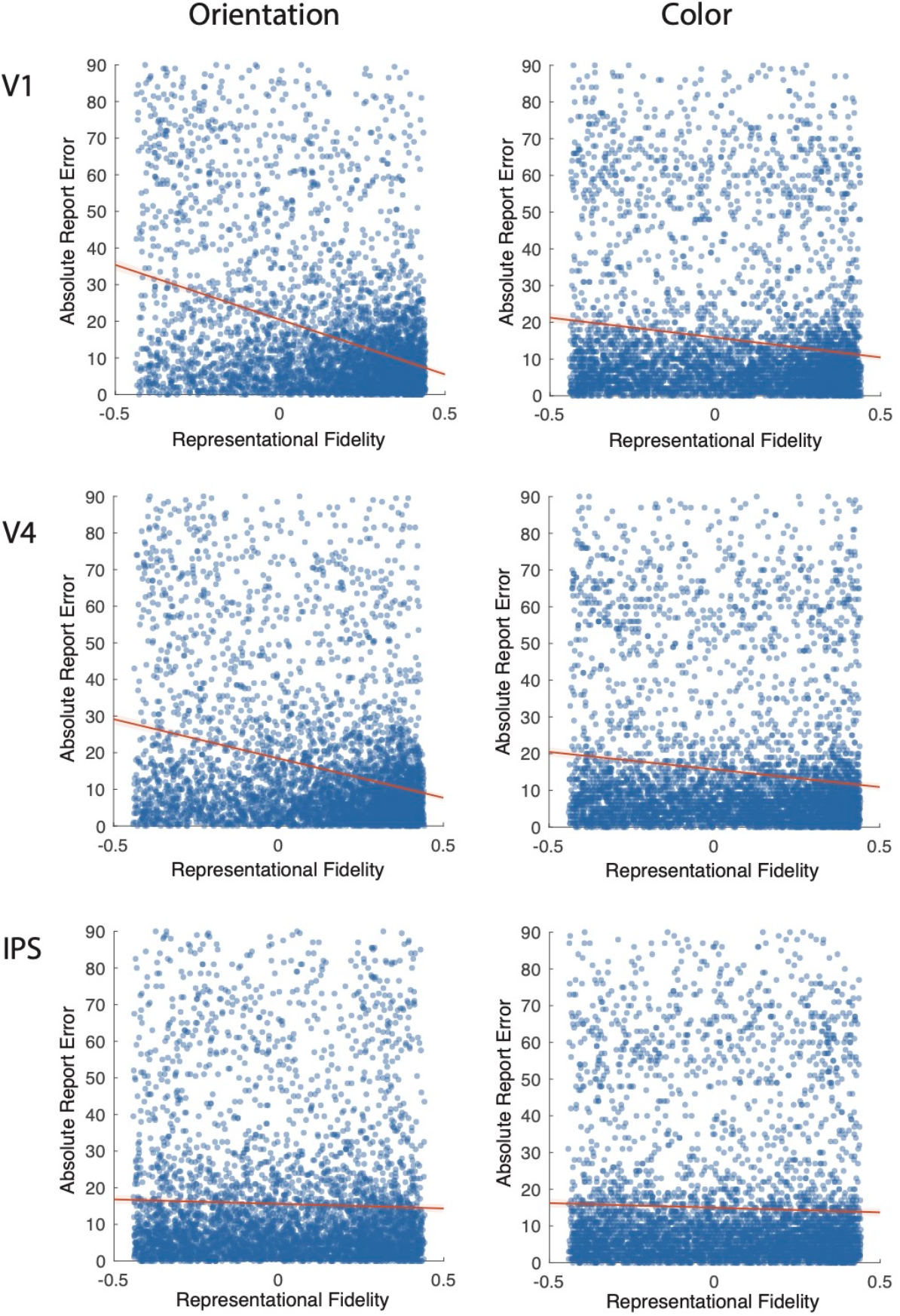
Neural reconstructions of task-relevant feature correlated with behavioral report. Trial-level Pearson’s correlation between RF of task-relevant feature and absolute report error. In all priori ROIs, we observed a negative correlation between the reconstruction quality and behavioral error: the better the reconstruction qualities, the smaller behavioral errors.

### Can we also reconstruct the *task-irrelevant* feature of the attended items?

#### A priori ROIs

Following the similar approach as above, we also tested whether the *task-irrelevant* (unreported) feature of the attended item could be reconstructed in early visual ROIs and IPS (Figure 6). In hold trials, the reconstruction fidelity of the unattended feature in early visual ROIs were not better than chance (RFs<0.004, ps>0.319). Similarly, in shift trials, we could not reliably reconstruct unattended feature in early visual ROIs (RFs<0.006, ps>0.076). We also calculated the reconstruction fidelity in anatomically defined IPS. No reliable orientation reconstruction was achieved in color reporting blocks (RFs<0.002, ps>0.639). We observed a significant reconstruction fidelity in hold trials when unattended color was reconstructed but the pattern was not replicated in shift trials (Hold Color: RF=0.014, p=0.037; Shift Color: RF=0.003, p=0.656). The significant reconstruction of Hold colors in IPS is intriguing but given the lack of significant patterns in other areas and conditions, it is hard to know for now whether this is reliable or simply due to chance.

**Figure 6.**
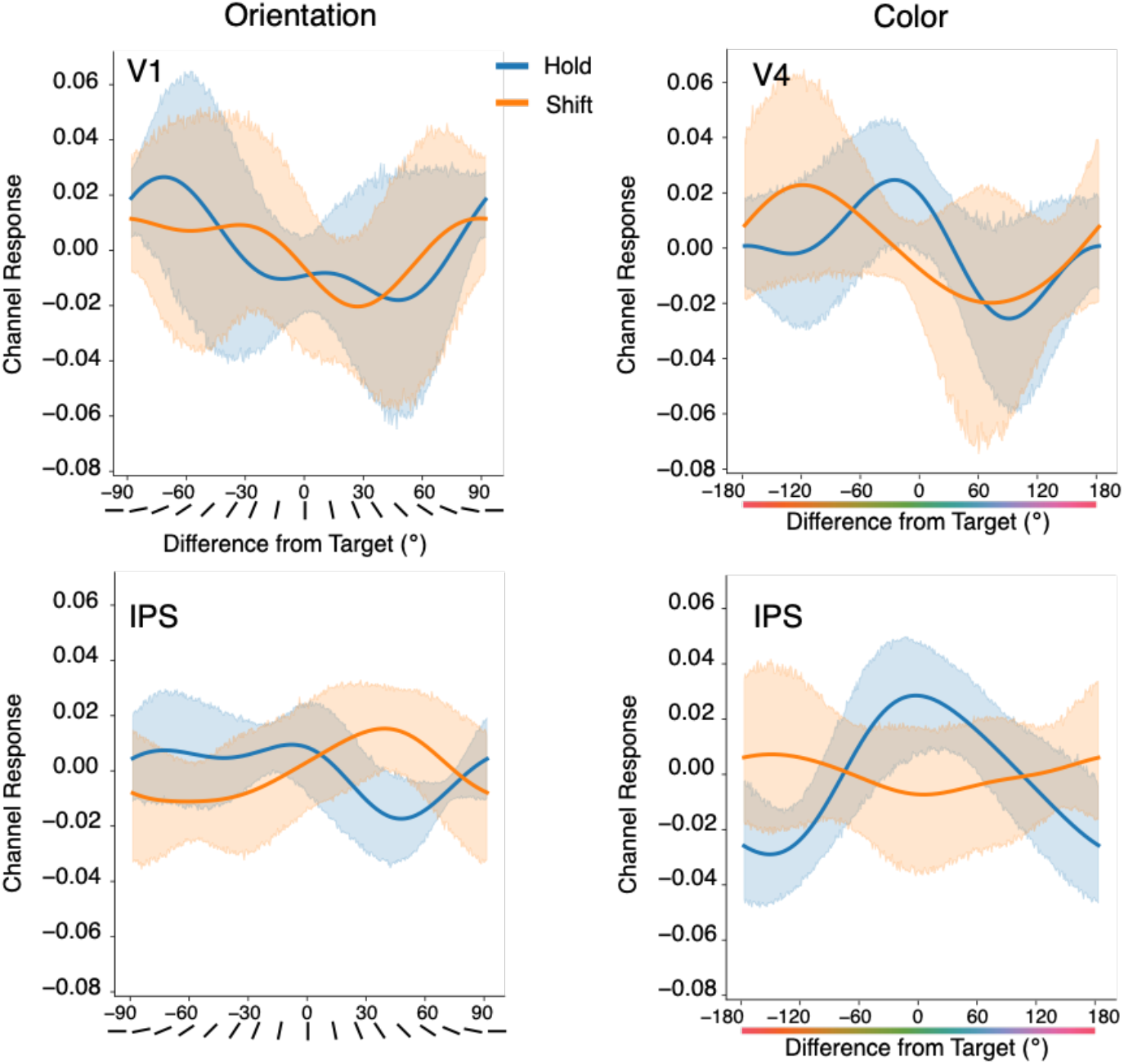
Neural reconstructions of task-irrelevant features of the attended item in priori ROIs. We did not observe significant reconstructions of task-irrelevant features in all condition and all ROIs except color reconstructions of hold trials in IPS.

#### Exploratory analyses in other occipito-parietal ROIs

We next did exploratory analyses to test whether task-irrelevant features could be decoded from other brain regions. First, we explored several additional anatomically defined occipito-parietal ROIs (Figure 7). For task-relevant features, we observed reliable feature reconstructions in most of our partial and occipital ROIs except for angular gyrus for hold trials. However, for task-irrelevant features, we did not find any significant reconstructions across these ROIs.

**Figure 7.**
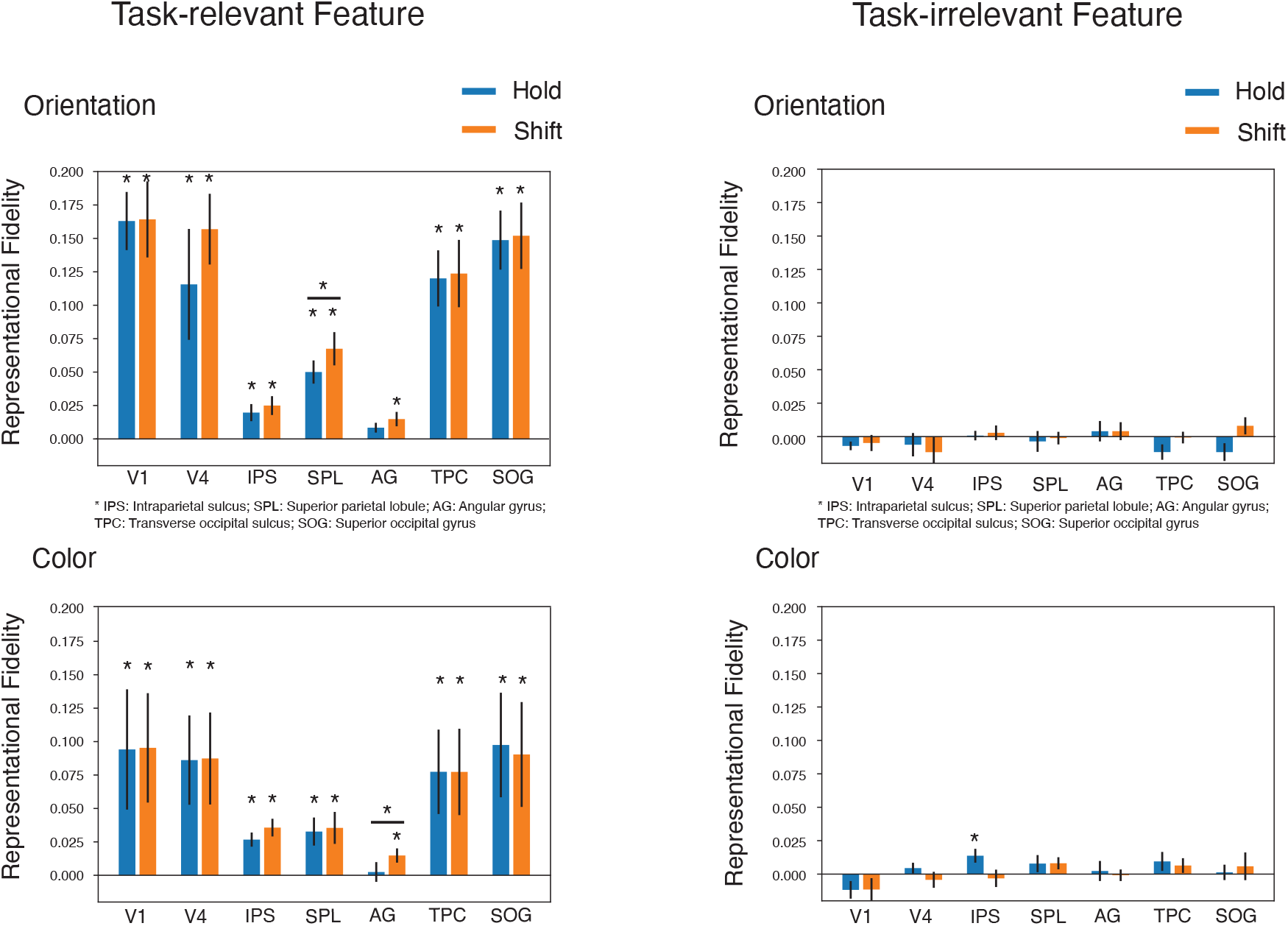
Neural reconstructions of task-relevant and task-irrelevant features of attended item in occipito-parietal ROIs. We observed significant reconstructions in all occipito-parietal ROIs for task-relevant features except angular gyrus for hold trials. However, we did not find any significant reconstruction in these ROIs for task-irrelevant features.

### Searchlight

Because we did not find any reliable reconstruction of the unattended features in the a priori ROIs, we next conducted a searchlight analysis to further explore whether any brain areas might possibly represent the task-irrelevant features. A similar analysis approach as for the ROI-based analysis was used for each spherical searchlight to generate a representational fidelity map in parietal and occipital regions. For comparison, representation fidelity maps were created for both attended features and unattended features. Because we did not find significant differences between hold and shift trials in feature reconstructions, we combined hold and shift trials to increase the statistical power.

The representation fidelity maps for attended features showed reliable feature reconstructions broadly in occipital and dorsal parietal regions (Figure 8). The reconstruction was strongest in occipital visual regions and gradually became weaker towards parietal regions. In contrast, the representation fidelity maps for unattended features revealed much weaker effects, but there were some significant clusters (Figure 8). For those trials that color was the task-irrelevant feature, the strongest color reconstruction was achieved in left posterior parietal cortex. For task-irrelevant orientation, we found a small significant cluster in right parietal cortex. The results suggest that there might be areas of parietal cortex which can reliably represent the task-irrelevant features, but further research would be needed to follow up on this exploratory result.

**Figure 8.**
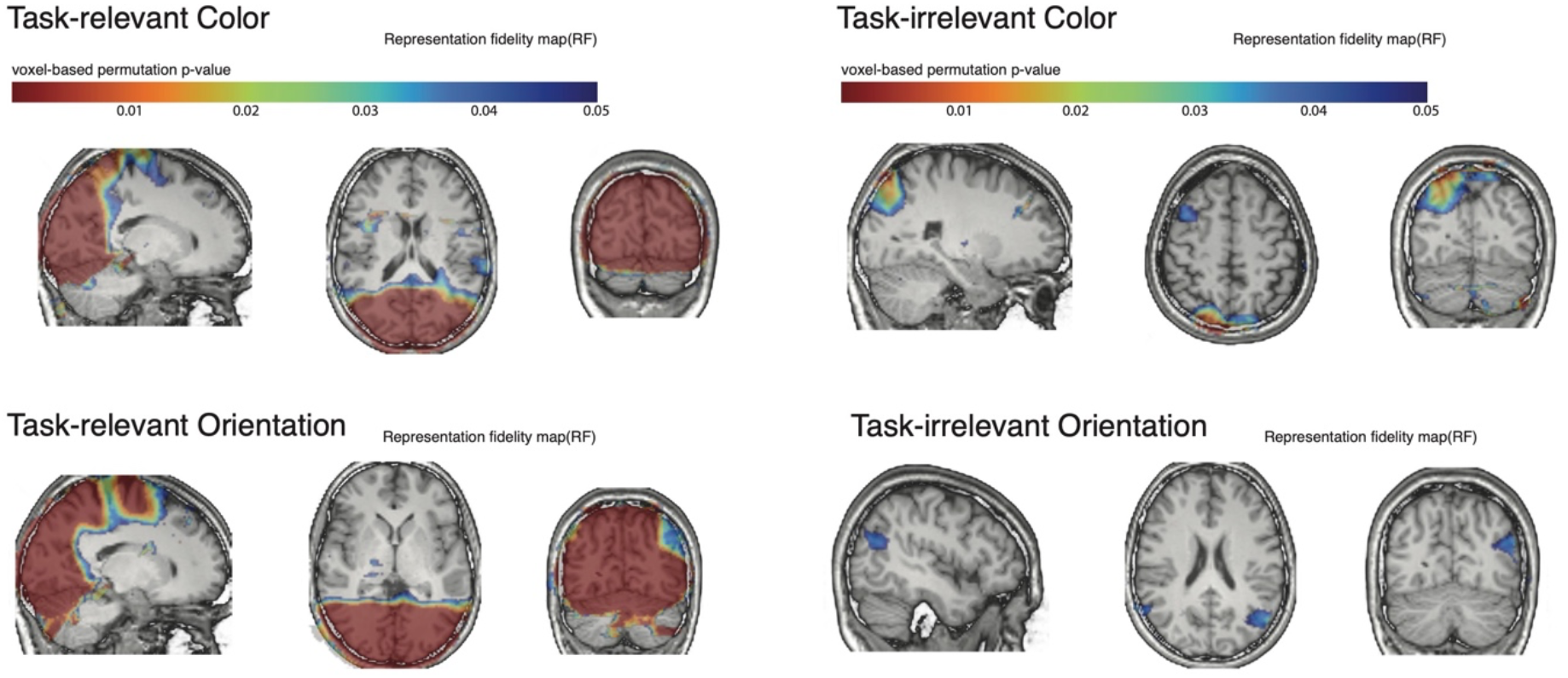
Searchlight maps of neural reconstructions for task-relevant features and task-irrelevant features. Maps are colored according to the permutation-tested significance of the RF value (without cluster correction); voxels with p-values greater than 0.05 are not shown. We observed reliable reconstructions for task-relevant features broadly in occipital and dorsal parietal regions but not for task-irrelevant features.

## Discussion

In the current study, we tested whether we could reconstruct the task-relevant and task-irrelevant features of the attended object in a fMRI task. We also asked how the reconstruction is affected by unstable spatial attention (dynamic shifts), and how the neural reconstructions link to behavioral performance. We successfully reconstructed the task-relevant features in a variety of visual sensory regions and parietal regions. The quality of the task-relevant feature reconstructions was correlated with the behavioral reports. We found that we could reconstruct the task-relevant feature of the item being spatially attended, even when spatial attention incorrectly selected the wrong object (swap errors). We also found that the quality of neural reconstructions (reconstruction fidelity) correlated with behavioral error on a trial-by-trial basis. Prior studies have demonstrated success at reconstructing task-relevant features from fMRI activity (e.g. Ester et al., 2015), but to our knowledge this is the first demonstration in a complex display with more than two multi-feature objects.

However, there are a few limitations of the current design that may affect the interpretation and robustness of these task-relevant reconstructions. First, participants made fewer behavioral swap errors than we had predicted based on our previous behavioral studies using a similar paradigm (Chen et al., 2019). The lack of big behavioral errors may be caused by extensive practice effects because participants practiced the task in two prescreening sessions and completed at least 4 hours of the task in the fMRI scanner. This reduced behavioral variability gave us less power to robustly explore trial-by-trial correlations between brain and behavior. The second limitation is more critical. As noted earlier, we decided to use a short memory delay after the stimulus presentation to increase the likelihood of detecting the task-irrelevant reconstructions. But this meant that the stimulus array and response period were temporally adjacent, such that the fMRI signal from the stimulus array may be contaminated by the signal from the response period. Thus it is possible that we reconstructed the participants’ response instead of the attended feature during stimulus perception. We took some measures in both the design and ROI selection (and additional control experiments; data not shown) to limit the extent of this confound, but we could not fully eliminate it. Thus, the task-relevant reconstructions should be interpreted with caution. That said, the more novel, main goal of this study was to test whether we could reconstruct the *task-irrelevant* features of the spatially attended item. The confound described above does not affect these task-irrelevant reconstructions, because participants were not asked to report those features. Thus, we focus primarily on those results.

We did not find robust evidence of reconstructing task-irrelevant features. We were unable to reliably reconstruct either task-irrelevant color or task-irrelevant orientation in our a priori early visual and IPS regions. Exploratory analyses found some intriguing clusters of significant task-irrelevant reconstruction in parts of parietal cortex, though this finding requires more study before claiming a solid conclusion.

There has been a long debate about the fate of task-irrelevant features (Bocincova & Johnson, 2019; Serences et al., 2009; Woodman & Vogel, 2008; Yu & Shim, 2017). Because of the very nature of the question, it is difficult to directly measure task-irrelevant feature representations with behavioral approaches. Previous behavioral attempts have reported mixed results on whether task-irrelevant features are processed or encoded (Marshall & Bays, 2013; Swan et al., 2016; Shin & Ma, 2016; Woodman & Vogel, 2008). Many studies provided evidence that even though task-irrelevant features may not be processed to the same level as the task-relevant features, they can still affect participants’ behavior (Marshall & Bays, 2013; Swan et al., 2016). One surprising single-trial test found that task-irrelevant features could be retrieved but with a much lower precision compared with task-relevant features (Shin & Ma, 2016). Our results suggest that task-irrelevant features could not be reconstructed from early visual regions using a linear encoding model like IEM. Thus, the behavioral influences of task-relevant features may need to be explained by something other than direct encoding of the task-irrelevant features in the same form as the task-relevant features.

Some previous studies in the working memory literature have similarly reported no evidence of active representation of task-irrelevant features in visual cortex (Yu & Shim, 2017). But because the representation of task-irrelevant features could be short-lived, the previous null results could be due to reconstructing from neural activity during the long delay before the memory response: i.e., the task-irrelevant feature could be processed initially but later dropped out during the memory delay. One advantage of our design (for the task-irrelevant representations) is that we minimized the memory delay after the stimulus display by presenting the response screen immediately after the mask disappeared. Therefore, a failure to reconstruct the task-irrelevant features in our study is hard to explain by the rapid dropout of the processing signal.

Studies on object-based attention suggest when attention is drawn towards one feature of the object, it facilitates the processing of the other features within in the same object (Duncan, 1984; Egly et al., 1994; O’Craven et al., 1999). Our results did not directly support that prediction. One possible reason is that participants may not group our two features (color and orientation) into one cohesive object. Xu (2010) used colored shapes (e.g. red square) as the objects, with color as task-relevant feature and shape as task-irrelevant feature. It is more natural in real life to group a red-square as a single object while a colored grating is extremely rare in our daily life. That might explain why shape in Xu (2010) was automatically processed as task-irrelevant features while color/orientation may not activate as the same level as shapes in our design.

Another possibility is that our task may encourage participants to strategically suppress the task-irrelevant features. Because color runs and orientation runs were blocked, it was easy to maintain a highly selective attentional filter to the task-relevant features and more efficiently ignore the task-irrelevant feature. One future design to consider would be to intermix the color trials and orientation trials within blocks, such that participants have to set the attentional filter every trial. In this alternative design, participants would still know which feature to report before each trial, therefore the unattended feature remains completely irrelevant in the current trial, but it could be task-relevant in the next trial. Participants are still not encouraged to remember the task-irrelevant features, but it is possible it may be harder to filter out the irrelevant features.

A final possibility is that the task-irrelevant features are represented in different forms compared with task-relevant features. For example, some studies suggested that the unattended contents in working memory are stored in an activity-silent form (Stokes, 2015). The attended features are maintained actively, and the contents could be decodable from population-level brain patterns (e.g., fMRI or EEG data). Temporarily irrelevant features are maintained in the activity-silent form and could be reactivated if they become relevant later. Although in current study the task-irrelevant features would not be tested later so participants had no incentive to reactivate them, it is still possible that that information was encoded in some “silent” form but cannot be reconstructed using linear models. That might explain why a study using univariate analyses could detect an overall signal increase when more task-irrelevant features were presented (Xu, 2010). Directly decoding/reconstructing task-irrelevant features may require further studies to build complex models to explicitly reflect the representational structure of how brain encodes those features.

In summary, in the current study we explored how the brain represents task-irrelevant features in a visual perception task. Although we did not find any evidence of significant reconstruction of task-irrelevant features in visual or parietal cortex, our results shed light on how the brain may represent task-relevant and irrelevant features differently and probes the extent to which spatial attention may act as a “glue” to bind task-relevant and task-irrelevant features.

